# Wheat - black-grass competition is a non-zero-sum game influenced by root growth

**DOI:** 10.1101/2025.06.02.657396

**Authors:** Jed Clark, Lynn Tatnell, Tom Bennett

## Abstract

Interactions with neighbouring plants have profound effects on plant growth, especially in crop-weed interactions, which can cause major crop-losses in agricultural systems. Black-grass (*Alopecurus myosuroides*) is the most problematic weed for UK agriculture, causing dramatic yield losses in winter wheat, but the basis of its competitive advantage is not clear. We aimed to fundamentally reappraise the nature of wheat – black-grass competition. Here, we show that black-grass is slow to establish, and requires long periods to demonstrate any competitive advantage over wheat. We show that black-grass grows significantly faster under winter conditions than wheat, and has much more vigorous root growth, collectively suggesting that prolonged root growth over winter is key to black-grass’s competitive advantage over wheat. We tested the competitiveness of a range of wheat and barley germplasm against black-grass in three distinct environmental set-ups, and show that competitiveness under controlled conditions reflects competitiveness under field conditions. In controlled conditions, we identified significant variation in crop varietal competitiveness against black-grass, in both ability to suppress black-grass growth, and to continue to grow in the presence of black-grass (tolerance). We further identified that this competition is not a zero-sum game, with loss in crop biomass not necessarily equal to gain in black-grass biomass. Finally, we found that the black-grass suppression, but not tolerance, is correlated with crop root growth, supporting our root growth hypothesis for black-grass competition. Overall, our results suggest that breeding for increased root growth and/or winter growth rate can create more competitive wheat cultivars for grassweed suppression.

## INTRODUCTION

Plants are active organisms, sensing and adapting to their environments in order to survive and reproduce. The presence of neighbouring plants is an unavoidable environmental stress in the life of most plants. Neighbouring plants compete for mineral nutrients and water in the soil, and for light-capture above ground, and thus directly impact the growth of a given focal plant. As such plants have developed multiple mechanisms of sensing neighbouring plants, including by changes in the spectrum of reflected light, the presence of volatile organic chemicals, and the presence of root exudates (reviewed in Bilas et al, 2021). The detection and response to these stimuli result in profound changes in shoot and root growth in focal plants, although the exact nature of the growth responses is still a matter of uncertainty. They can appear to the observer to be competitive, neutral or cooperative, but the extent to which plants are deliberately following such strategies is controversial, and there often are other explanations for observed changes in growth, such as confounding variables and inherent asymmetries between species (reviewed in Bilas et al, 2021). Understanding plant-plant interactions has significant implications for the growth of crop plants in agricultural settings (Becker et al., 2023). Crops are typically densely sown, and there each plant unavoidably impacts the growth of its neighbours. As such, it is very likely that crops have been bred for reduced individual growth, that allows all plants to attain similar sizes, and results in maximised field yield (at the cost of reduce individual yield) (Weiner et al., 2010). Maximising crop yields thus to some extent requires overcoming the downregulation of growth that typically occurs in neighbouring plants, while still maintaining quasi-cooperative behaviour among neighbours (Wheeldon et al., 2022). Interactions between crop and weed plants add another dimension to this puzzle, with ecological simplification to monoculture fields providing perfect conditions for the proliferation of weeds (Cusworth and Lorimer, 2023; Gandy, 2023). Weeds represent a particular problem to crops precisely because crops have been bred for reduced individual growth (Weiner, 2017); weeds therefore often have a dramatic competitive advantage over individual crop plants, and can rapidly decimate the yield of a crop field (Mennan and Isik, 2004).

Black-grass (*Alopecurus myosuroides* Huds.) is an agricultural grass weed across Western Europe (CABI, 2008). Black-grass is a highly competitive weed, which produces very high seed-sets, making it impossible to completely remove from the seed-bank once fields are infested (Lutman et al, 2013). For winter wheat plantings, black-grass competition causes significant yield reductions and is the most problematic weed species for winter wheat in the UK, a problem that has rapidly escalated in the last 40 years (Moss et al, 2007). Management of black-grass has traditionally relied heavily on herbicide application. After ploughing or dressing of empty fields in the autumn, black-grass seeds will germinate and can be ‘sprayed off’ before the crop is drilled into the field (Bastiaans, Paolini and Baumann, 2008). However, the problem of black-grass infestation has been heightened by the fact that many populations of black-grass are now herbicide resistant, due to the over reliance on selective herbicides (Moss, 2007; Moss, 2017). Other management approaches therefore accept the presence of black-grass, but aim to reduce its advantage over wheat. September drilling of winter wheat is preferable, since it increases yields and is a better period for using machinery within fields, but early-sown crops are particularly susceptible to black-grass infestation. Delayed sowing into October can decrease black-grass presence, but reduces the growth period for wheat, and means sowing takes place at a less easy period for cultivation (Lutman et al., 2013). Spring plantings of wheat, are less affected by black-grass, with reductions of between 78-96% black-grass relative to September sowing (Lutman et al., 2013). Some crops, such as winter barley, are also significantly less impacted by black-grass than wheat, but the reason for this is currently unclear (Cook and Roche, 2018).

Within wheats, competitiveness against black-grass is cultivar dependant, with more competitive lines reducing black-grass mean head counts between 22-30% compared to less competitive varieties (Lutman et al, 2013). Therefore, it has been suggested that competitive cultivars could be used as a cultural control method to naturally suppress black-grass (Andrew et al, 2015). This integrated weed management (IWM) method could be used alongside reduced herbicide input (Harwood, 2020) to reduce the impact of black-grass. IWM however has its limitations. Highly variable results at the individual field level limit their uptake (Lutman et al, 2014) with failing control methods having practical and economic consequences for the farmer. The use of competitive cultivars however, has been highlighted as a cost-effective and compared to other methods, and a labour free approach to IWM (Andrew et al, 2015). Unfortunately, there is no quick and easy way of assessing the competitiveness of a cultivar (Andrews, 2016). Identifying competitive cultivars requires an understanding of wheat-black-grass competition, which is currently lacking. Previous studies have surmised possible above-ground mechanisms through which wheat – black-grass competition may occur. Traits such as increased crop height (Wicks et al. 1986; Huel & Hucl 1996; Lemerle et al. 1996) and aspects of canopy architecture (Champion et al., 1998) have been linked with black-grass suppression or maintenance of crop yields in the presence of black-grass (black-grass tolerance). The impact of tiller production on competitive ability has seen contrasting results with some reports showing increased competitive ability in higher-tillering wheat (Lemerle et al. 1996; Hucl 1998; Korres & Froud-Williams 2002) and some only showing weak correlation (Wicks et al. 1986; Champion et al. 1998). More recently, tallness and early heading were associated with black-grass tolerance and suppression, but it was highlighted that other factors must also be responsible for competitive ability in wheat (Mason, Goonewardene and Spaner, 2008). Andrews, (2016) conversely suggested that found prostrate growth habit may increase competitiveness. Due to the difficulties in investigating root systems under field conditions, no studies have looked at below-ground parameters as a potential explanation for the outcome of competition between wheat and black-grass.

The aim of this study was to fundamentally re-appraise the nature of competition between wheat and black-grass, by studying competition under controlled conditions at a much higher level of detail. In our experiments, we were able to look at root growth as potential explanation for competition, and we tested the hypothesis that differences in elemental root growth resulted in the competitive advantage of black-grass over wheat. By screening different wheat and barley cultivars for competitive ability against black-grass, we were able to link these abilities, and in particular black-grass suppression to root growth. We conclude that root growth is likely a key indicator of competitive winter wheat and barley varieties, and a suitable target for trying to breed new competitive cultivars.

## MATERIALS & METHODS

### Plant materials

Wheat lines, both winter and spring, were used for plant growth experiments. These lines consisted of current elite, and landrace varieties. Barley lines used in experiments included spring, winter, and hybrid barleys. Seed from a black-grass population was obtained from ADAS, Boxworth field C. Seed from the same population was used for all black-grass experiments. All lines were obtained either from current university stocks or from breeders. Unless otherwise stated experiments in winter conditions used winter wheat variety Claire and winter barley Bordeaux, whilst experiments in spring conditions used spring wheat Mulika and spring barely Bowman.

### Plant growth conditions

Soil-based experiments used Petersfield Potting Supreme No. 2 compost. Experiments under ‘spring’ conditions were grown in glasshouses at 20°C with supplementary LED lighting with an average intensity of 250 μmolm^−2^s^−1^ on 16-hour day and 8-hour night cycle. Experiments under ‘winter’ conditions were carried out in Weiss growth cabinets at a 10°C, with supplemental LED lighting with an average intensity of 198 μmolm^−2^s^−1^ on an 8-hour day 16-hour night cycle. For germination and early growth assessments plants were grown in a Walk-In cabinet at 20°C, with supplemental LED lighting with an average light intensity of 310 μmolm^−2^s^−1^ on a 16-hour day 8-hour night cycle.

### Black-grass seed viability

To ensure seed viability for experiments all black-grass seed was pre-germinated prior to sowing in soil. Seeds were pre-germinated in Petri-dishes on filter-paper saturated with water and grown at room temperature. Once germinated black-grass seedlings of a similar size were selected for sowing.

### Determining germination characteristics of black-grass and elite winter wheats

Seed was sown into Petri-dishes each containing filter paper saturated with water and placed within the Walk-In cabinet. Seeds were assessed for germination (defined as the first emergence of either the root or coleoptile from the seed) every 24 hours. Once roots/coleoptile were visible their length was recorded every 24 hours up to 14 days post germination.

### Determining short term growth rate of black-grass and elite winter wheats

Pre-germinated seed was transferred to 100ml P24 pots containing soil. The pots were placed into glasshouse conditions for two or four weeks. After two or four weeks respectively, the shoots were harvested at the soil surface. Harvested shoots were dried in an oven at 60 °C for seven days after which shoot biomass was recorded using a balance.

### Hydroponic plant growth and root analysis

Ungerminated seeds were sown in 100ml P24 pots consisting of a 1:1 mix of sand and perlite. Seeds were grown in glasshouse conditions for 14 days after which similar sized plants were identified and selected. Selected plants were carefully removed from their pots, ensuring the root system remains intact. The roots were then washed in water to remove remaining sand/perlite. The root-shoot junction was encased by a foam bung and placed within a modified, open-ended falcon tube (cut to 4-5cm in length) with shoots protruding from one end and roots the other. For each plant, a 1L hydroponic pot was filled with water. An air pump and air stone connected by tubing were used to supplying oxygen and agitation. The falcon tube containing the plant was placed within a hole in the hydroponic pot lid, orientating plant roots into the hydroponic pot. Pots were placed into either glasshouses or Weiss cabinets depending on the individual experiment. The water within the hydroponic pots was topped up as required. Every two weeks the whole pot was drained and replaced with fresh water. Upon set-up and subsequent addition of fresh water, 15ml of Arabidopsis thaliana salts (ATS) solution (Wilson et al, 1990) was added to each pot. After six months the plant shoots and roots were harvested. The shoots were cut at their base where it meets the top of the falcon tube. The roots were cut from their top at the base of the falcon tube. Any plant material remaining within the foam bung was discarded. The roots and shoots were placed into a drying oven for seven days after which dry root and shoot biomass was recorded along with tiller number.

### Rhizobox plant growth and root analysis

Rhizoboxes were made from Corning 245mm X 245mm square plates. The base of the plates were modified in two ways, 1) Small holes for water uptake were created on the base edge of the plate using a hot metal skewer, 2) A rectangular hole 5×2cm was cut into the top edge of the plate to allow vertical shoots growth. Soil was sieved to remove large particles, before being placed into the plates ensuring tight compaction. The soil was then watered. A singular seed was sown into the soil approximately 1-2 cm from the top edge of the plate in its centre. The lid was then attached to the plate. The plate was then covered with an opaque plastic cover (also sporting a hole for the shoot) and placed at 45^0^C in a metal rack ensuring the base of the plate was facing downwards. The rack was the placed within a water filled tray ensuring the base of the plates are submerged. Rhizoboxes were placed in glasshouses and grown for 10 weeks. Plants were harvested after 10 weeks. Shoots were cut at their base at the top of the rhizobox. Shoots were placed within seed bags and then into a drying oven for seven days after which dry shoot biomass was measured using a balance.

To produce root images, the top and base of the rhizobox plates were periodically scanned. Images were assessed using ImageJ software. Percentage root cover was recorded, this assumed that roots are white, and soil is black. Percentage root cover was calculated through white pixel assessment. Using ImageJ, each pixel within each image was graded for intensity on a scale of 0-255, where 255 = completely white and 0 = completely black with the numbers in-between being the range from white to black. The mean pixel intensity therefore calculates the amount of white in the whole image, this value is relative on a scale of 0-255, which is converted to a percentage. This gives a value for percentage white pixel cover and therefore a value for percentage root cover. For our images, the polygon tool was used to draw around the desired part of the image, ensuring to avoid areas of the plate not consisting solely of soil or root, as this would affect the result. White pixel cover was then recorded within the selected area. For these experiments the image contrast was set at 25/299 and was kept constant across all images.

### Controlled environment wheat competition screen

For glasshouse experiments, 2L pots were filled with soil. A single crop seed was sown at 1-2cm depth in the centre of the pot. For the competitive setup only, six pre-germinated black-grass seed were sown at 1-2cm depth at equidistance in a hexagonal formation around the centre crop. Crop tiller number was recorded every seven days. After three months crop and black-grass shoots were harvested at the soil surface. The shoots were dried in an oven at 60 °C for seven days after which shoot biomass of all individual crop plants was recorded. For black-grass, the biomass of all black-grass plants within the same pot was weighed together as one. For winter experiments the method was the same except plants were placed into the Weiss cabinet and harvested after six months.

### Container trials

Container trials took place from October 2021 to July 2022 at ADAS’s Boxworth site in Cambridge. The trials consisted of 23 different treatments each with four replicates giving a total of 92 pots organised in a fully random split block design. Pots used were 20L circular pots, filled with sterilised loam mix (Rothamsted ‘weed mix’ - Sterilised kettering loam and Lime free grit 3-6mm in a 4:1 ratio plus 2kg/m^3^ Osmacote) to a depth of 2 cm below the rim. The pots were placed on an uncovered hard-standing area and watered well using an overhead watering system. The area was fenced to prevent predation from ground animals (rabbits etc.). In all treatments except black-grass only treatments, pots were sown with 10 crop seed of corresponding variety, at a depth of 1-2cm. Seed was positioned evenly across the pot surface avoiding sowing within 15mm of the pot edge. Crop seed was thinned to six plants per pot after emergence (four weeks post-sowing) whilst at the leaf stage of emergence (GS11-12), ensuring equal spread of remaining crop plants. In treatments 8-23 upon sowing, an uncontrolled number of black-grass seed was scattered onto the pot surface then covered with a fine layer of soil to an even depth of 1cm. These pots would later be thinned after emergence (four weeks post sowing) to achieve the desired black-grass densities (10 or 20 plants per pot), ensuring BG remaining after thinning were equally spread across the pot. In treatments 8-15 the pots were thinned to 10 black-grass plants per pot, treatments 16-23 were thinned to 20 black-grass plants per pot. The pots were left outside throughout the winter growing season. Pots were watered as required. No chemical or biological treatments were given to the plants. Pots were checked for other weeds such as annual meadow grass and weeded if necessary. First plant assessments were carried out in March 2022. These assessments consisted of tiller counts on both crop and black-grass. The tiller number of each crop plant and each black-grass plant in every pot was recorded. Further assessments were carried out in May 2022. This consisted of counting the total number of black-grass ears present in each pot. In July 2022 the pots were harvested by hand. For each pot the plant shoots were cut at soil level and all above ground biomass was collected. Crop and black-grass biomass was separated by hand, final wet shoot weight was recorded for both for each pot.

Unfortunately, barley seed was heavily predated by birds prior to harvest therefore recorded barley (Bordeaux and SY Kingsbarn) biomass’ will be lower than the maximum they achieved. All wheat varieties and black-grass showed no signs of predation. Small amounts of biomass may have been lost at harvest and transport as well as human error in separating crop from black-grass biomass.

**Supplementary Table 1:** Experimental set-ups for container trials, showing combination of wheat (W), barley (B) and black-grass (BG) plants in each treatment.

| Treatment | Density |
| --- | --- |
| 1. Kerrin | 6W |
| 2. Elation | 6W |
| 3. Barrel | 6W |
| 4. W1190637 | 6W |
| 5. W1190488 | 6W |
| 6. Bordeaux | 6B |
| 7. SY Kingsbarn | 6B |
| 8. Kerrin + black-grass (low) | 6W + 10BG |
| 9. Elation + black-grass (low) | 6W + 10BG |
| 10. Barrel + black-grass (low) | 6W + 10BG |
| 11. W1190637 + black-grass (low) | 6W + 10BG |
| 12. W1190488 + black-grass (low) | 6W + 10BG |
| 13. Bordeaux + black-grass (low) | 6B + 10BG |
| 14. SY Kingsbarn + black-grass (low) | 6B + 10BG |
| 15. Black-grass (low) | 10BG |
| 16. Kerrin + black-grass (high) | 6W + 20BG |
| 17. Elation + black-grass (high) | 6W + 20BG |
| 18. Barrel + black-grass (high) | 6W + 20BG |
| 19. W1190637 + black-grass (high) | 6W + 20BG |
| 20. W1190488 + black-grass (high) | 6W + 20BG |
| 21. Bordeaux + black-grass (high) | 6W + 20BG |
| 22. SY Kingsbarn + black-grass (high) | 6W + 20BG |
| 23. Black-grass (high) | 20BG |

### Field trials

The field trials took place from October 2022 to July 2023 at ADAS’s Boxworth site in Cambridge. A field with an existing black-grass population was selected. The trial consisted of eight different treatments, seven crop treatments and one black-grass only control. Each treatment had four replicates giving a total of 32 plots. One treatment (W1190637) only had enough seed for three replicates therefore the fourth plot was covered with Extase. Plots were 5m x 2m and crop seed were sown at seed rates of 325seed/m2 using an Oyjord drill at a depth of 4cm. Black-grass was also hand sown to all plots to ensure complete coverage over the whole trial area. Crop plots were randomised within the trial area, black-grass only plots were placed at the end of the trial area to ease drilling. Four days post drilling, glyphosate (Samurai 4l/ha) pre-emergence herbicide was applied to the trial area.

First assessments were taken in April 2023 at crop GS30-31. In each plot, black-grass plants were counted within a 0.5m^2^ quadrat which was placed randomly within the plot avoiding the outermost rows. Average crop height (three plants) and crop tiller number (three plants) per plot was recorded for each plot. Further assessments were carried out in June 2023. In each plot black-grass ears were counted within a 0.5m^2^ quadrat which was placed randomly within the plot, again avoiding the outermost rows, this was done by holding the quadrat above the plants and counting the number of ears that were within the quadrat square. Crop height (three random plants) was also recorded for each plot. In July 2023 the plots were harvested. Grab samples were taken from each plot. The grab samples were collected as follows; two 0.5m^2^ quadrats were randomly placed in each plot. All plants that had their shoots originating within that quadrat were cut at soil level, bundled together, and collected. The two grab samples from each plot, were then combined to form a single biomass sample for each plot and transported to the on-site lab. Crop and black-grass biomass were then separated, as was any biomass from any other weed species present, this unwanted biomass was discarded. Once separated, crop and black-grass wet shoot biomass was recorded. Primary crop tiller number (tillers with ears) was also recorded.

### Statistical analysis

Sample size and statistical output for each experiment is stated in the figure legends. Data was tested for normality using the Shapiro-Wilk test to determine the appropriate statistical tests for each dataset.

## RESULTS

### Black-grass is slow to establish and weedy early in life

One possible explanation for the competitiveness of black-grass against wheat is that black-grass seedlings gain a ‘head-start’ on wheat by prodigious germination or vigorous seedling growth. Given the small size of black-grass seed, we did not deem this especially likely, but we wanted to test this explanation nevertheless. We therefore examined several seedling vigour parameters in black-grass and three elite winter wheats (Elation, Barrel and Kerrin). We measured: 1) time from imbibition to germination (defined as the time to first visible emergence of either root or coleoptile), 2) time to reach a root length of 4cm, 3) time to reach a coleoptile length of 2cm; 4) shoot biomass after two weeks and 5) shoot biomass after four weeks. Black-grass seed took on average four days longer to germinate than all three elite wheats (Figure 1A). Coleoptile growth took black-grass significantly longer than the wheats, taking 2-3 times longer to reach 2cm (Figure 1B), while black-grass roots growth took significantly longer than wheat to reach 4cm (Figure 1C). Shoot biomass of black-grass after both two and four weeks was much smaller than any of the wheats (Figure 1D,E). Thus, the data show that black-grass is slow to germinate and establish compared to wheat, and unlikely to gain any early competitive advantage over wheat.

**Figure 1:**
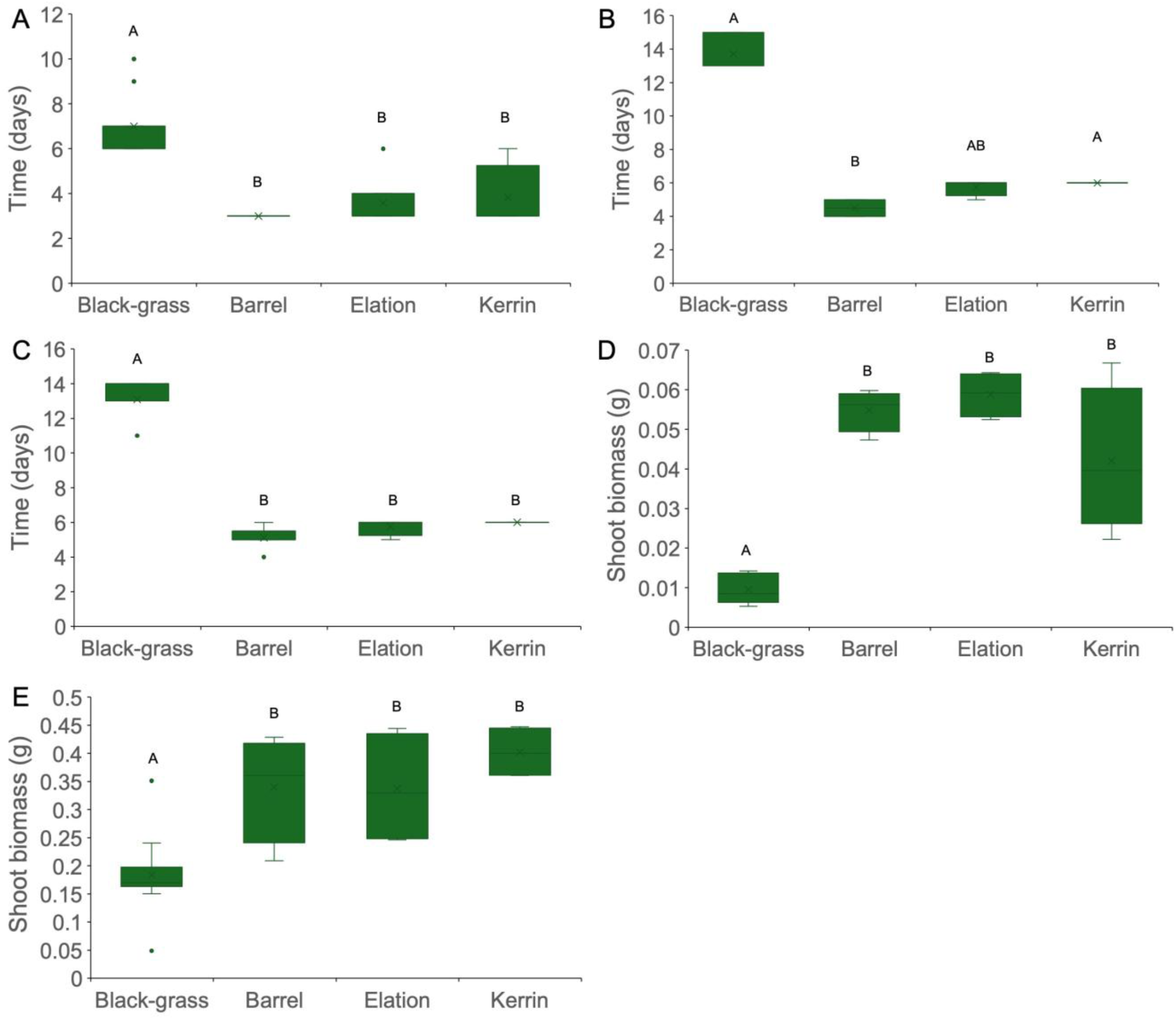
Black-grass seedlings are slow to establish in comparison to wheat. **A-E)** Boxplots comparing the early growth of black-grass and three elite wheats (Barrel, Elation and Kerrin). Panels show (**A**) time taken to germinate (n=6-11), (**B**) the time taken to produce a root of 4cm in length (n=4-8), (**C**) the time taken to produce a coleoptile of 2cm in length (n=4-11), (**D**) seedling shoot biomass 2 weeks post germination (n=4-11) and (**E**) seedling shoot biomass 4 weeks post germination (n=4-12). Boxes indicate interquartile range, the midline indicates the median, the whiskers are the minimum and maximum values, the X within the box is the mean. Circles represent outliers. Different letters above the boxes indicate significant statistical differences between the groups, calculated for panels **A**, **B**, **C** and **E** by (Kruskal-Wallis, Dunn’s adjusted with Bonferroni correction, P<0.05) and for panel **D** by (ANOVA, Tukey’s HSD test, P<0.05).

### Black-grass competition is slow to develop

Despite being slow to establish, and weedy early in life, black-grass still eventually comes to outcompete winter wheat in the field. To identify when black-grass gains a competitive advantage, we conducted a competition experiment in both spring and winter conditions. Winter wheat (Claire) was grown in 2 litre pots either as solitary wheat plants, or sharing the pot with six black-grass plants. Crop tiller number was recorded throughout the experiment as a proxy for progressive shoot growth. Under ‘spring’ conditions (16 hours light, 20°C), it took 66 days to observe a significant difference between the tiller number of plants in the treatment group compared to the control group (Figure 2A). In more realistic ‘winter’ conditions (8 hours light, 10°C), a significant difference in tiller number between tiller number was first seen after 118 days (Figure 2B). Thus, even at a 6:1 ratio of black-grass to wheat, it takes a very long time for black-grass to have any noticeable impact on wheat shoot growth. This reflects agricultural experience, where black-grass is often invisible until spring. These data suggest that the sheer amount of time spent in the field might explain the greater competitiveness of black-grass in winter wheat plantings compared to spring wheat plantings.

**Figure 2:**
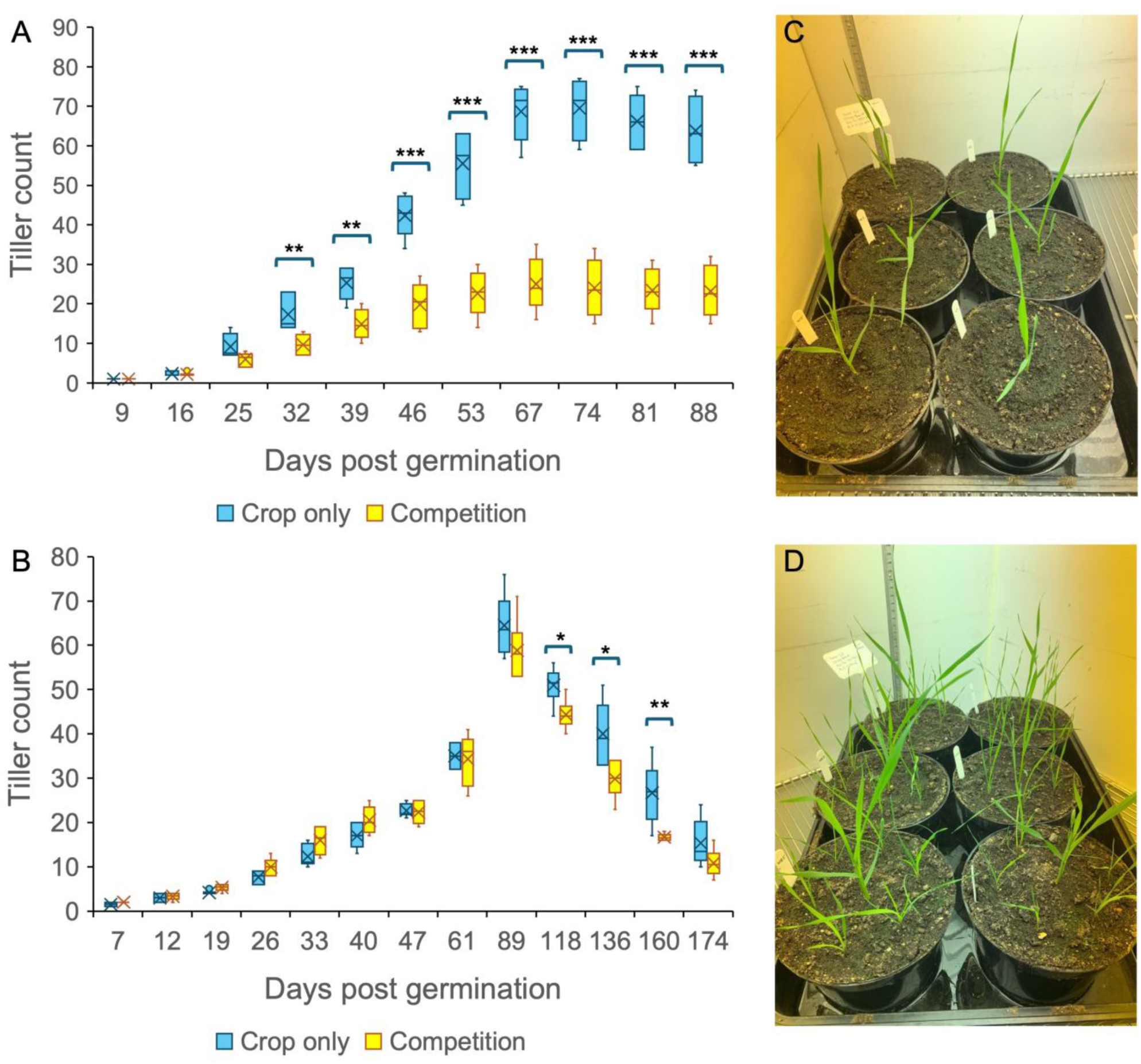
Black-grass competition is slow to develop. **A,B)** Boxplots showing crop tiller number over time post germination with and without black-grass competition for both (**A**) spring and (**B**) winter growth conditions. The box indicates the interquartile range, the midline indicates the median, the whiskers are the minimum and maximum values. X denotes the mean. Asterisks denote statistically significant differences between groups (Independent samples t-test/Mann-Whitney U-test, P<0.05) (*<0.05, **<0.01, ***<0.001), n=6. **C,D)** Images showing the experimental set up without (**C**) and with (**D**) competition from blackgrass.

### Black-grass has greater winter growth rate than wheat, especially in the root system

While time can explain the difference between winter and spring wheat, it cannot explain the difference in black-grass competitiveness between winter wheat and winter barley. Since elite barleys generally have better developed root systems than elite wheats, we hypothesised that the difference in root growth rate between wheat and barley might explain these observations. Furthermore, we hypothesised that black-grass might have greater root growth than wheat, and that this is the source of its competitive ability. To test this idea, we grew winter wheat (Claire), winter barley (Bordeaux) and black-grass in a hydroponic system in winter conditions, to allow easy measurement of root growth. At the end of the experiment, root and shoot biomass were measured. In this experiment, both barley and black-grass produced much larger root systems than wheat (Figure 3E-G), and also significantly larger shoot systems, with a lower shoot:root ratio (Figure 3A-C). While blackgrass produced a larger proportion of root biomass than barley, barley had an equal root biomass to black-grass because of its overall faster growth. These data are therefore consistent with root growth as an explanation for the competitive advantage of black-grass over wheat, but not barley.

**Figure 3:**
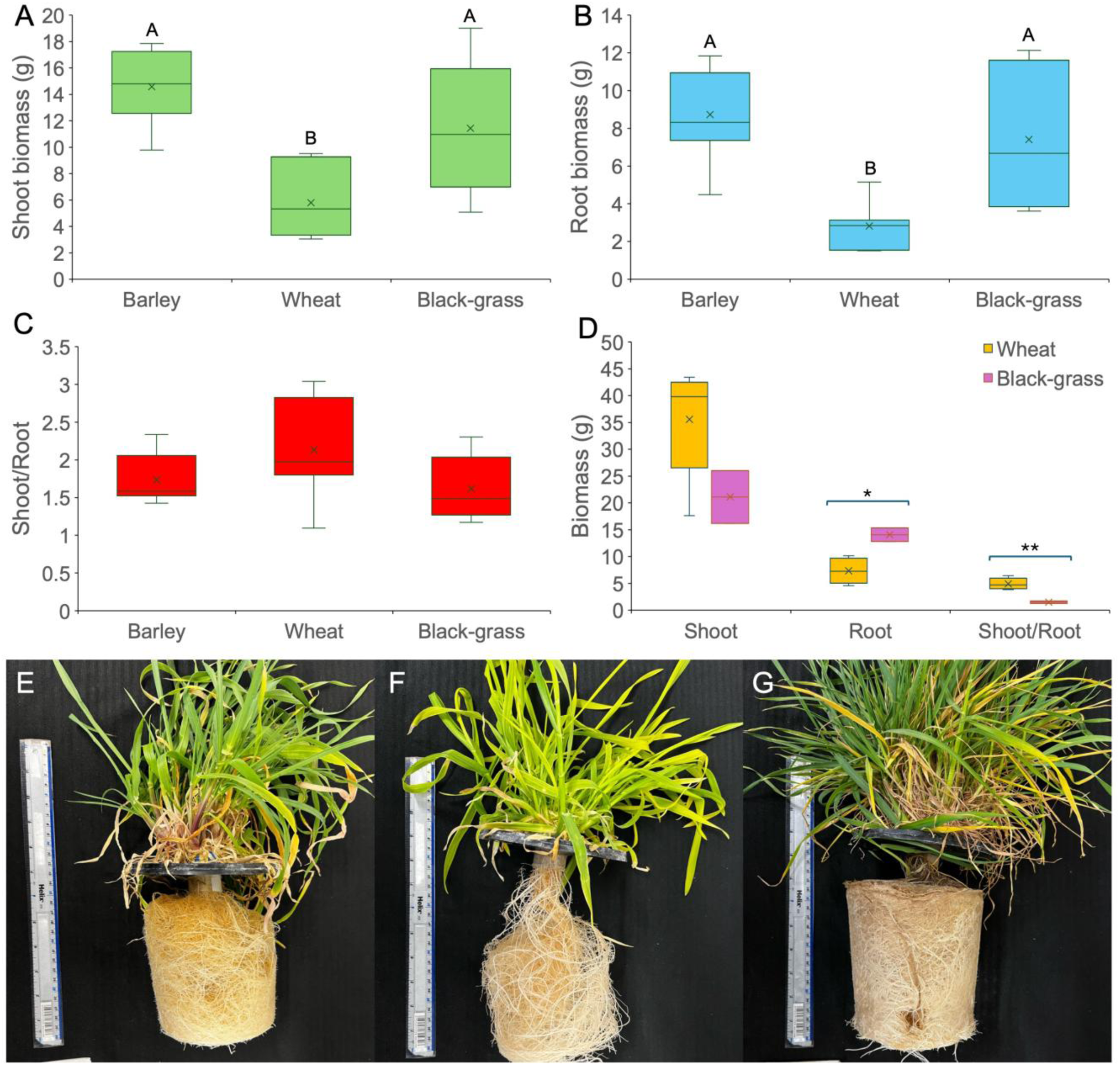
Black-grass grows more strongly than wheat in winter conditions. **A-C)** Boxplots showing the growth of Barley, Wheat, and Black-grass in a hydroponic system in winter conditions with (A) showing the final dry shoot biomass, (**B**) the final dry root biomass and (**C**) the final dry shoot to root biomass ratio. Different letters above the boxes indicate significant statistical differences between the groups calculated for (**A** and **B**) by (ANOVA, Tukey HSD test, P<0.001), n=6-8. **D)** Boxplot comparing wheat and black-grass growth in spring conditions. The box indicates the interquartile range, the midline indicates the median, the whiskers are the minimum and maximum values, the X within the box is the mean. For (**D**) Asterisks denote significant statistical differences between the groups (Independent samples t-test *=P<0.05, **=P<0.01), n=2-6. **E-G)** Images of root growth in hydroponics in winter conditions of barley (**E**), wheat (**F**) and black-grass (**G**) after 167 days of growth.

We were surprised by the low growth of wheat in this system, given its vigorous growth in spring conditions. We therefore conducted another hydroponic experiment to compare wheat and black-grass growth under spring and winter conditions. We found that in spring conditions, winter wheat (Claire) had much faster and stronger growth than black-grass, resulting in a much higher shoot biomass (although still a smaller root system)(Figure 3D). Conversely, in winter conditions, black-grass had stronger growth in both compartments than wheat (Figure 3A-C), with a much smaller reduction in growth between spring and winter conditions than observed in in wheat (compare Figure 3A/B and D).

Collectively, our data suggest that black-grass has a competitive advantage over winter wheat in agricultural settings because it allocates a larger proportion of biomass to its roots, because it has stronger growth in winter conditions, and because it has sufficient time to overcome its poor establishment. These three factors allow black-grass to dominate below-ground resources and space by the spring, at which point it becomes a suddenly-noticeable issue within the field. Conversely, black-grass is less competitive with spring wheat due to the reduced time in which to build a large root system, and less competitive than winter barley because barley plants grow faster than wheat, and have larger root systems.

### Root growth properties vary amongst wheat and barley varieties

To test this ‘root growth hypothesis’, we decided to look at variation between different wheat and barley lines for root growth. We reasoned that if lines with larger root systems were also more competitive against black-grass, our hypothesis would be supported. We therefore used two methods to assess root growth, which give slightly different data. Firstly, we used ‘rhizoboxes’ to assess root growth, which allow the visualization of roots grown within soil, although in this method the root system is compressed into a thin into a quasi-2D growth space (see methods). The rhizoboxes were repeatedly imaged, and the percentage coverage by roots at each time point was used as a proxy for root growth. We screened 24 lines using this approach, which included two wheat landraces from the Watkins collection (W1190637 and W1190488), four barleys (Tardis, Feeris, Bordeaux and SY Kingsbarn), black-grass and 17 elite wheat lines. Due to space considerations, rhizoboxes experiments were carried out in spring conditions, and were grown in separate batches. The elite winter wheat line Kerrin was used as an internal control, included within each batch, allowing direct comparisons between batches using the percentage of the root growth relative to the internal Kerrin control. The four Kerrin samples showed very similar root growth between batches, so the absolute data for all lines is shown on Figure 4A. Under these conditions, we observed reasonable variation in root growth between wheat lines from ∼30% to ∼47% root cover. Consistent with our observations in Figure 3, barley lines showed very strong root growth, although black-grass itself showed lower root growth in these conditions.

**Figure 4:**
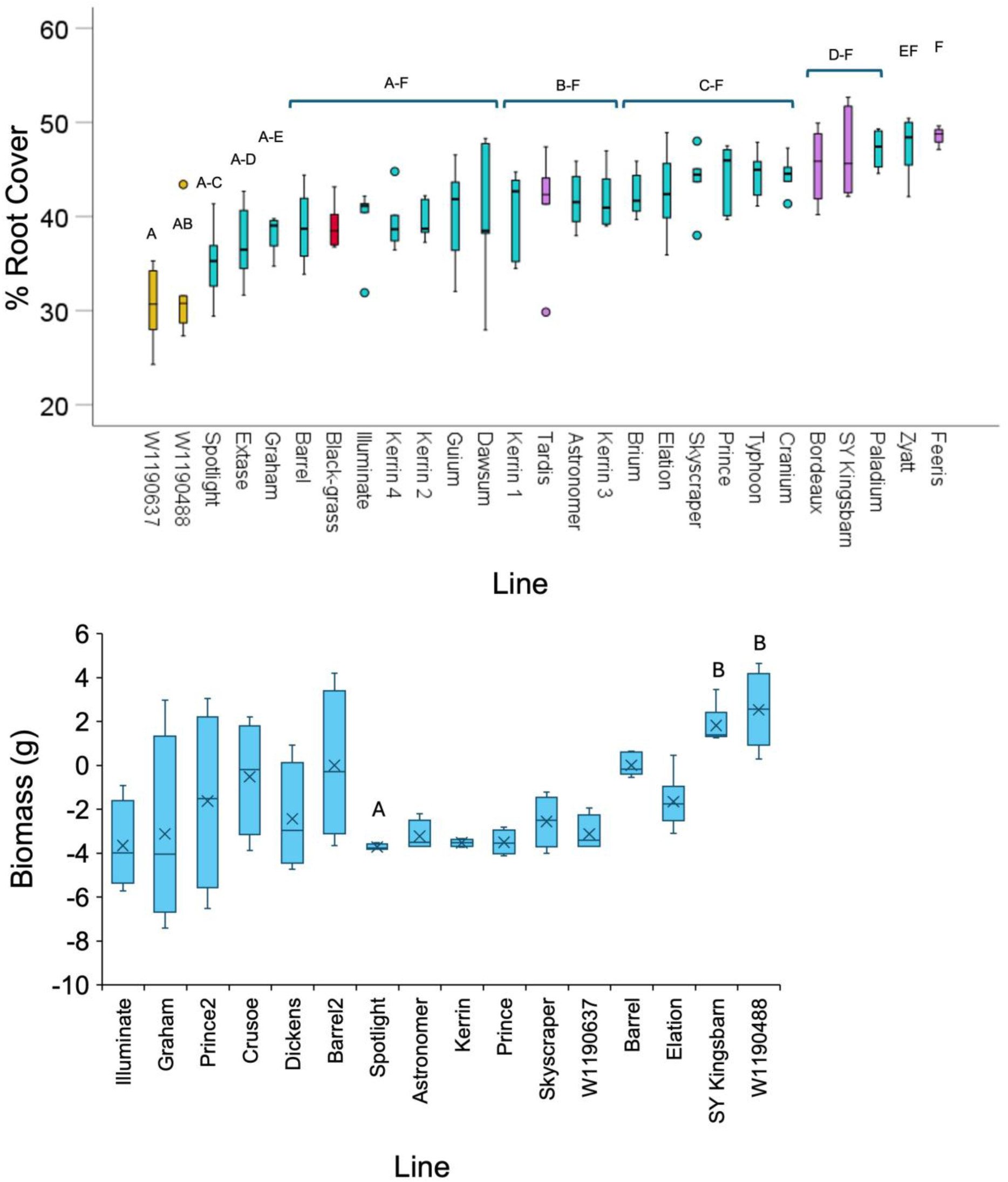
Root growth varies between wheat, barley and black-grass lines. **A)** Box-plot showing the percentage root cover of the total rhizobox surface for each crop variety. red = black-grass, yellow = wheat landrace, pink = barley, blue = elite wheat **B)** Boxplots showing hydroponic crop root biomass for 16 crop lines in both batch one and batch two. All data is normalised to the Barrel control within the corresponding batch. The box indicates the interquartile range, the midline indicates the median, the whiskers are the minimum and maximum values. Circles represent outliers. Samples with the same letter are not statistically different to each other; for (**A**) Kruskal-Wallis, Bonferroni correction, p<0.05., n=5, for (**B, C**) (ANOVA, Tukey’s HSD, P<0.05), n=4.

A sub-set of 14 lines (One barley, two wheat landraces, 11 elite wheats) were then selected and grown in a hydroponic growth system in winter conditions (Figure 4B), to gain a more accurate picture of root growth under the relevant growth conditions in which wheat – black-grass competition is established. The use of the hydroponic system already described allowed direct quantification of final total root biomass. Again, due to space constraints, plants were grown in separate batches, with the winter wheat Barrel as an internal control. The two batches were quite different in terms of absolute root growth, with one batch showing more extensive variation than the other. In Figure 4B, the root biomass of each line is therefore plotted relative to its internal Barrel control. The results show important dissimilarities to the growth under spring conditions in rhizoboxes. Barrel showed the strongest growth of the elite wheat lines, which was not the case under spring conditions. The other elite wheats were rather similar to each other, which is comparable to the spring conditions, as is the strong growth of SY Kingsbarn in both conditions. However, one of the landrace wheats (W1190488) showed the strongest root growth in winter conditions, despite having the weakest growth in spring conditions. Overall, while there was a correlation between root growth in the two conditions, it is clear that root growth in winter conditions cannot be confidently predicted from growth in spring conditions.

### Wheat and barley varieties consistently vary in competitiveness against black-grass in multiple environments

To assess whether lines with different root growth vary in competitiveness against black-grass, we selected seven of the previously tested crop varieties – three elite winter wheat cultivars (Barrel, Elation, Kerrin), two landrace winter wheat cultivars (W1190637 and W1190488), one winter barley (Bordeaux) and one hybrid winter barley (SY Kingsbarn). We hypothesised that the lines with greater root growth would show greater competitiveness against black-grass, across multiple environments. To test this idea, we used three different experimental set ups. Firstly, we grew solitary wheat plants with six black-grass neighbours (as per Figure 2) under controlled glasshouse conditions. Secondly, we grew 10 wheat plants in 20 litre containers, with either no black-grass, 10 black-grass plants (low density) or 20 black-grass plants (high density). These microcosms were grown outside on a hard-standing in Cambridgeshire, UK, in the 2021/22 growing season. Thus, although soil volume, water availability and plant densities were controlled, the plants were subject to the natural weather conditions of that season. Finally, we grew the crop varieties in a field trial in Cambridgeshire in the 2022/23 growing season, in 5 x 2m plots. Crops were sown at a density of 325 seed per m^2^, and then over-sown with black-grass seed. For each experiment, we also included a black-grass-only treatment (no sown crop). For all three experiments, final crop biomass (per pot, per container or per plot) we measured, along with the final black-grass biomass in the same areas.

Although there is some variation between the experiments, they generally show a consistent pattern of results. The black-grass-only controls have the highest black-grass biomass in each condition, and provide a useful reference for the degree of black-grass suppression by the crop varieties. SY Kingsbarn showed the greatest suppression of black-grass in both container and field trials and the second highest in the glasshouse trial, while Bordeaux showed the greatest suppression in the glasshouse, and second greatest in the field trial (Figure 5A-C). The lack of black-grass suppression by Bordeaux in the container trial therefore seems highly anomalous (Figure 5B). Consistent with agricultural practice, the barley varieties were therefore highly competitive against black-grass.

**Figure 5:**
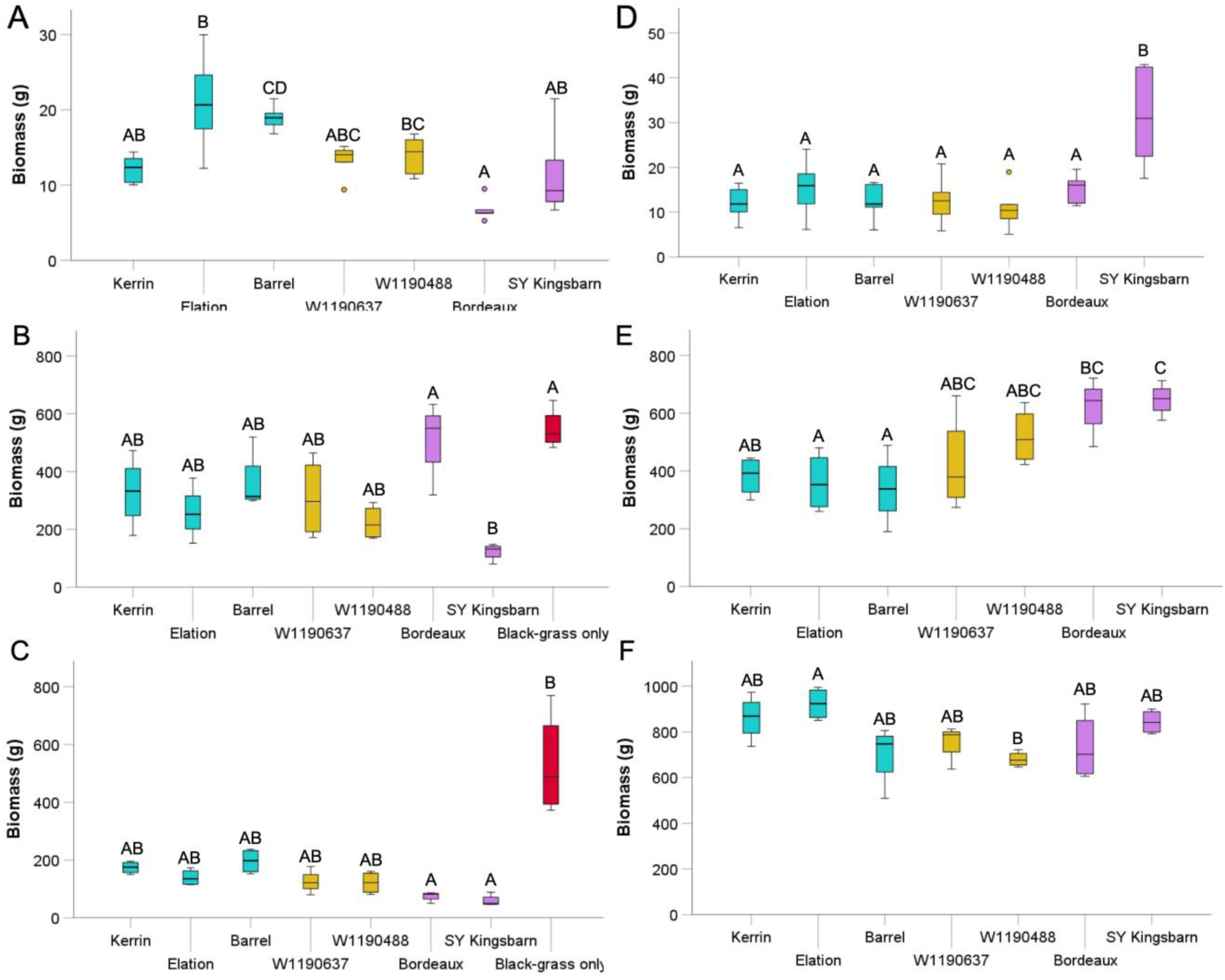
Competitive ability of crop lines is consistent across environments. **A-F)** Boxplots showing summary of competitive ability of seven crop lines across three different environments. Boxplots show black-grass shoot biomass (**A**, **B** and **C**) and crop shoot biomass (**D**, **E** and **F**) in controlled glasshouse conditions (**A** and **D**), container trials (**B** and **E**) and field trials (**C** and **F**). Different letters denote significant statistical differences between groups denoted for (**A**, **B**, **C** and **D**) by (Kruskal-Wallis, Dunn’s with Bonferroni correction, P<0.05) and for (**E** and **F**) by (ANOVA, Tukey’s HSD test, P<0.05). Colours represent crop type, blue = Elite wheat, yellow = Landrace wheat, pink = Barley, red = black-grass only. Controlled growth conditions (n=6), container trials (n=4) and field trials (n=4).

By contrast, the three elite wheats were generally very comparable to each other in terms of black-grass suppression. The two least-suppressing lines in each experiment were elite wheats; Barrel and Elation in the glasshouse, and Barrel and Kerrin in both the container and field trials, with Elation the third least suppressing line in the field trial (Figure 5A-C). Kerrin performed surprisingly well in the controlled environment, and Elation the same in the container trial (Figure 5A,B). Nevertheless, the elite winter wheats were generally less effective at supressing black-grass across all experiments. The two landrace wheats were also generally very comparable to each other, and were intermediate between the elite wheats and the barleys in each experiment (Figure 5A-C).

The crop biomass measurements are somewhat more difficult to interpret, as there are inherent differences in the size of the crop plants when grown in isolation. In the glasshouse and container trials, both barleys had large biomasses consistent both with black-grass suppression, but also the vigorous growth of barley (Figure 5D,E). The wheats were generally very similar in the glasshouse (Figure 5D), but in the container trials, the two landraces had the largest biomass, consistent with their suppression of black-grass (Figure 5E). In the field trial however, both Elation and Kerrin produced the highest biomass, exceeding even SY Kingsbarn, while Bordeaux, Barrel and the two landraces had lower biomasses. Thus, even though Kerrin and Elation had high black-grass burdens, they still grew exceptionally well under field conditions, perhaps consistent with their specific breeding under these conditions.

### Competitive ability against black-grass varies between crop genotypes

Taken together, our results show that the competitiveness of varieties against black-grass is generally comparable across different environments. However, the results did not show a clear relationship with the root growth observed in either rhizoboxes or hydroponics. To try and get a better handle on this, we decided to assess a broader range of crop varieties under glasshouse conditions, and more carefully measure the loss of crop biomass between solitary plants and those grown with six black-grass neighbours, and the corresponding increase in black-grass biomass under the same conditions. We could not grow all varieties at the same time, and we observed significant batch-to-batch variety in terms of absolute growth. However, by measuring 1) the percentage loss in crop biomass between control, 2) the percentage gain in black-grass biomass and 3) the percentage change in the total plant biomass between the control and treatment, we could still directly compare varieties based on their relative performance. Using this system, we grew 26 different varieties, including the 24 shown in Figure 4A, with two additional elite winter wheats.

These experiments, showed stark differences in how well different crop lines are able to compete and retain crop shoot biomass when in competition with black-grass, with a range of 13-80% loss of biomass between different lines (Figure 6A). Consistent with other results, barleys generally do best in these experiments, and the two least-affected lines are both barleys. The seven least-affected lines only lost between 13-45% biomass, and interestingly included six of the lines discussed in Figure 5 (excepting Barrel). Thus, Elation and Kerrin actually represent some of the more black-grass-competitive elite wheats. The remaining lines, all but one of which are elite wheats, all lost between ∼60-80% of their biomass, consistent with the observed poor performance of elite wheats under field conditions.

**Figure 6:**
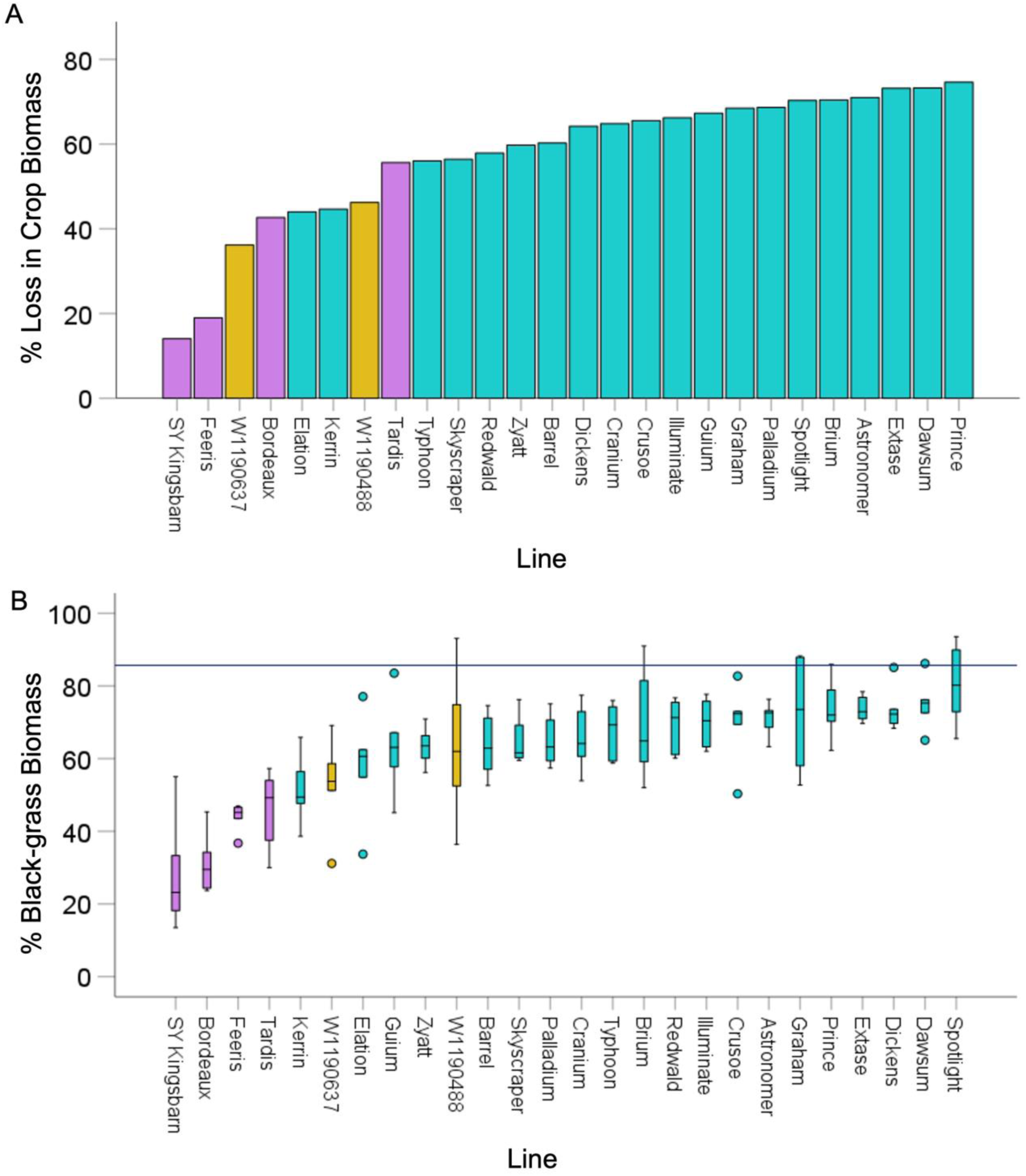
Crop lines vary in black-grass tolerance and suppression. (**A**) Bar graph showing average percentage loss in crop biomass in competition with black-grass, relative to crop only controls, n=6. Pink = barley, yellow = winter wheat landrace, blue = elite winter wheat. (**B**) Boxplot showing the percentage of the total pot biomass that is black-grass for different crop lines all in competition with black-grass. The box indicates the interquartile range, the midline indicates the median, the whiskers are the minimum and maximum values. Circles represent outliers, n=6. Blue line indicates the expected black-grass biomass in a pot if all plants grew to the same size and therefore the effect of competition being null. Bar colours represent crop type, blue = Elite wheat, yellow = Landrace wheat, pink = Barley.

These experiments also showed stark differences in the proportion of the total biomass that is black-grass, varying between 25% and 80% (Figure 6B). These increases were generally (but not absolutely) well-correlated with the loss in crop biomass. The best four blackgrass-suppressing lines were all barleys, and six of the seven best black-grass-suppressing lines were also in the seven least-affected lines in terms in crop biomass. Thus, there are considerable differences between the competitive ability of different crop lines against black-grass, which are significantly greater than those observed in Figure 5 described above.

### Wheat – black-grass competition is a non-zero-sum game

Interestingly, while there is a good correlation between loss in crop biomass and gain in black-grass biomass, the correlation is not absolute (Figure 7A). Thus, the gain in black-grass biomass is not necessarily equal to the loss in crop biomass. Indeed, for most lines, there is an overall gain in the total biomass in the pot when black-grass and crop is grown together (Figure 7B). However, some lines show a reduction in overall biomass when grown together (Figure 7B). These data strongly suggest that wheat-blackgrass competition is not a zero-sum game, and that the losses of one plant do not simply equal the gains the other. There are some clear general trends among this data. As previously shown, the least-affected crops are typically the best at suppressing black-grass (Figure 7A). However, in these lines, the suppression of black-grass is somewhat less effective than would be expected from the lack of change in crop biomass; black-grass gains significantly more than the loss in crop biomass, such that the system as a whole over-produces. The only exception here is Bordeaux, which suppresses black-grass strongly, but the system as a whole shows a loss of productivity. Conversely, in the most affected wheat lines, there is generally a tighter connection between loss of crop biomass and gain in black-grass biomass. Nevertheless, for some varieties the system as a whole still over-produces.

**Figure 7:**
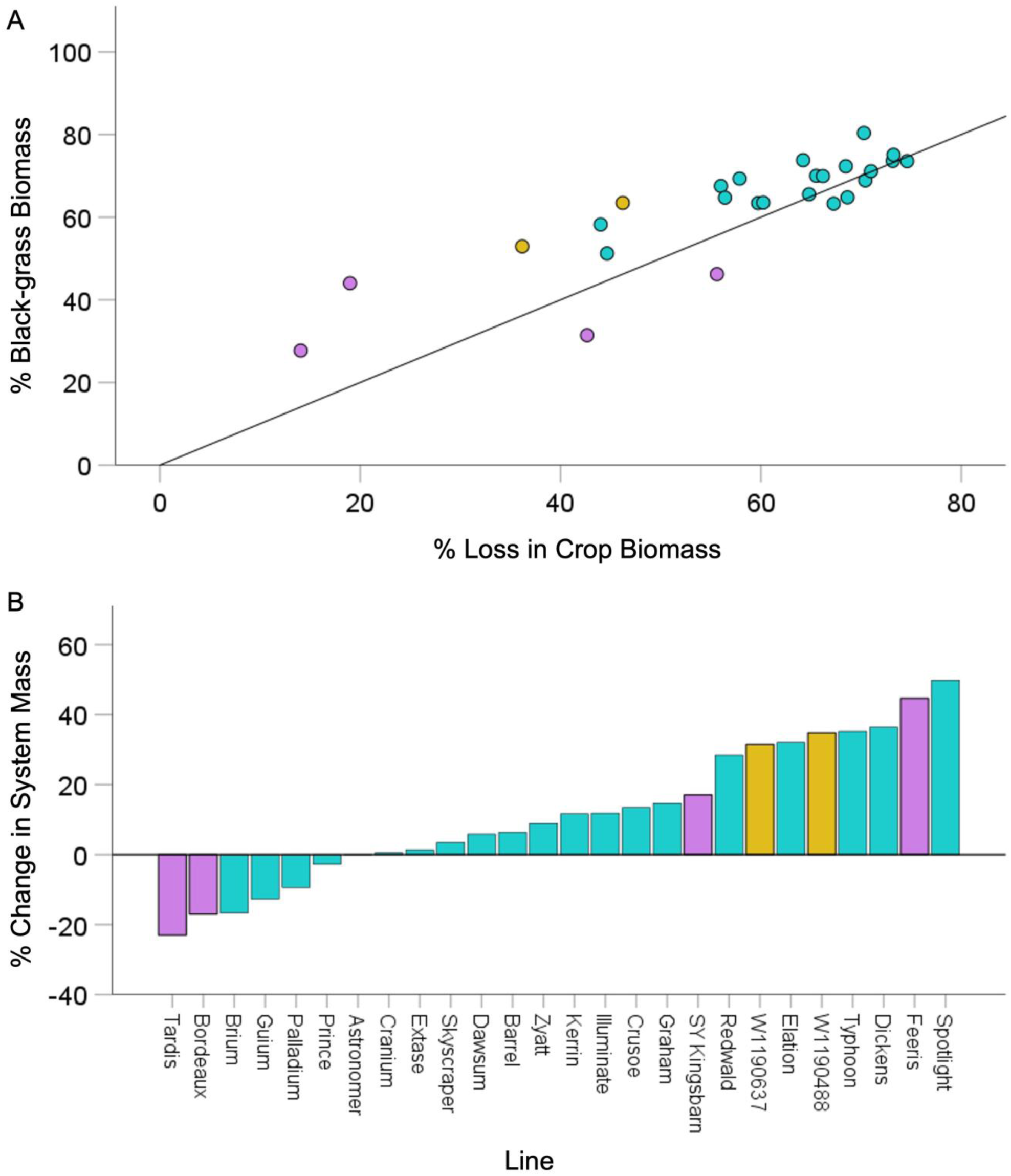
Crop and black-grass competition is not a zero-sum game. (**A**) Scatterplot showing the relationship between percentage total pot biomass that is black-grass biomass and percentage loss in crop biomass in competition with black-grass compared to the crop only controls. Points are averages. Line = y=x, representing a zero-sum relationship (**B**) Bar chart showing the average change in total pot biomass productivity from crop only pots to crop - black-grass competition pots. Zero-sum line denoted by Y=0. Colours represent different crop type, Red = Barley, Blue = Landrace wheat, Green = Elite wheat. (N=6). Colours represent crop type, pink = barley, yellow = winter wheat landrace, blue = elite winter wheat.

At a more nuanced level, the data also suggest that there are two different traits among crop varieties; the ability to continue growth in the presence of black-grass (‘black-grass tolerance’), and the ability to suppress black-grass growth (‘black-grass suppression’). While these are generally correlated, and might be inter-related, the data suggest they are not the same thing. For instance Bordeaux seems excellent at suppression (Figure 6A), but less good at toleration (Figure 6B), at least in these experiments.

### Black-grass suppression is likely related to root growth

Having defined the competitive ability of different varieties against black-grass, we then wanted to test whether our root growth hypothesis was correct, and whether this competitive ability relates to the root growth of crop varieties. To do this, we took the rhizobox root growth data (Figure 5A), normalised as a percentage of Kerrin, and plotted it against the percentage loss in crop biomass for each line (i.e. black-grass tolerance) (Figure 8A) and the percentage gain in black-grass biomass for each line (i.e. black-grass suppression) (Figure 8B). The results show no clear connection between black-grass tolerance and root growth (Figure 8A), but do show a clearer correlation for black-grass suppression (Figure 8B). This is particularly the case if the values for the two winter wheat landraces are removed from the data, which can be justified on the basis that their root growth in rhizoboxes was anomalously small compared to their growth in hydroponics (especially WW190488). If this is done, then there is a relatively good correlation between root growth and lack of black-grass growth in the dataset (Figure 8C). We therefore conclude that root growth likely plays an important role in the competitiveness of crop varieties against black-grass, by allowing them to monopolise space/resources in the rhizosphere earlier in the growing season, thereby suppressing the growth of later-developing black-grass plants.

**Figure 8:**
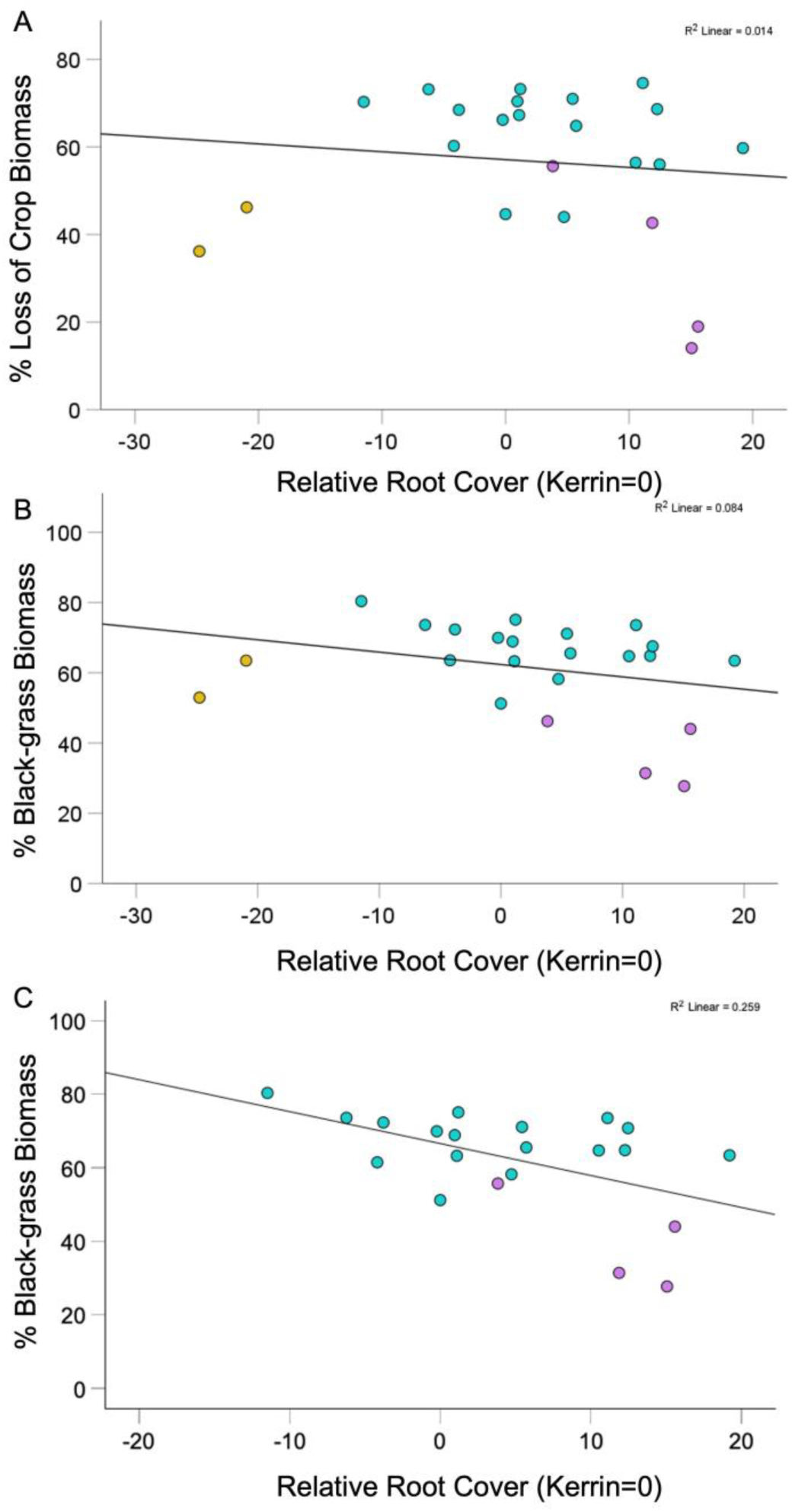
Black-grass suppression is correlated with crop root growth. Scatterplots showing the relationship between competitive ability and root growth, root growth assessed by % coverage of the rhizobox surface (**A**) percentage loss in crop biomass in competition with black-grass, relative to crop only controls, plotted against root growth in the rhizobox assay as a percentage of the internal Kerrin control sample (**B**) percentage of the total pot shoot biomass that is blackgrass, in crop/blackgrass competition samples, plotted against root growth in the rhizobox assay as a percentage of the internal Kerrin control sample (**C**) excluding winter wheat landraces, showing the relationship between the percentage of total pot biomass that is black-grass, plotted against root growth in the rhizobox assay as a percentage of the internal Kerrin control sample. Kerrin root growth =100%, represented as zero on the x-axis. Colours represent crop type, pink = barley, yellow = winter wheat landrace, blue = elite winter wheat. Trendlines represent linear lines-of-best-fit.

## DISCUSSION

### A new model for wheat – black-grass competition

The data presented here allows us to propose a new model for wheat – black-grass competition, and provide some explanation for why black-grass is much less problematic in spring wheat and winter barley plantings. Consistent with its wild and weedy nature, black-grass is slow to germinate and establish compared to wheat (Figure 1), and as such, black-grass takes a long time to grow to sufficient size to compete with neighbouring wheat seedings – even when at a 6-fold advantage (Figure 2). Thus, above all, black-grass only becomes a problem in winter plantings, because only there is there enough time for competition to establish. These findings are supported by Lutman et al., (2013) who found changing crop planting densities had no impact on black-grass presence in the autumn and spring but had a significant impact by the summer, showing the delayed nature of black-grass wheat competition. However, black-grass’s advantage in winter planting is redoubled by its faster elemental growth in colder, short-day conditions compared to wheat (Figure 3). In warmer, longer-day conditions, wheat grows faster than black-grass, further limiting the effects of black-grass in spring plantings.

The combination of longer available time and faster growth in winter conditions is sufficient for black-grass’s “hidden weapon” of massive root system growth (Figure 4) to take effect. Thus, over winter, black-grass wins an underground battle for space and nutrient capture, allowing to subsequently outgrow wheat above ground in spring. Our model provides an explanation for the apparently sudden appearance of black-grass in spring in fields that had appeared relatively unaffected. Our data suggest that winter barley is less susceptible to black-grass because it has faster growth than wheat in winter conditions, and faster elemental growth per se, allowing it to maintain the same pace of root system growth with black-grass. Previously, above ground traits have been linked with barleys increased competitiveness in comparison to wheat (Cook and Roche, 2018), but below-ground comparisons have not been made. Further work is required to definitively test our model, ideally by using by using high/low root growth variants of the same wheat cultivar to show that – all other factors being equal – root growth determines the outcome of competition.

Our data also provide some explanation for the dramatic expansion of black-grass as an agricultural problem in the UK over the last 40 years (Moss, 2017a). The geographical spread of black-grass within the UK is likely due to the movement of seed between fields on the wheels of agricultural machinery (Walsh, Newman and Powles., 2013), but the intensification of black-grass infestations within fields requires a different explanation. The spread of herbicide resistance contributes to this intensification (Hull et al., 2014), but two other factors also likely contribute. The first is warmer winters, due to the effects of global warming, which likely exaggerate the growth advantage of black-grass over winter wheat, while the second is the declining root system size of winter wheat as a result of crop breeding (White et al, 2012), which again exaggerates the inherent advantages of black-grass over wheat. This ‘perfect storm’ of factors has led to major crop losses in UK agriculture to black-grass infestations (Moss, 2010).

### Restoring the roots – a multiple win scenario for wheat breeding

A major trend in wheat breeding over the last 40 years has been an emphasis on ever-higher harvest indexes, with a greater and greater proportion of biomass allocated to the seeds (Reynolds, Rajaram and Sayre., 1999). This trend has greatly reduced the canopy size of elite wheat shoot systems, with particularly noticeable reductions in flag-leaf size, for instance. It is likely the same breeding trend accounts for the progressive reduction in elite wheat root system size of the same time period (White et al, 2012), which has been facilitated by the expansion of high-input agricultural practice, meaning crops can be successfully grown with smaller root systems. However, our results indicate that this reduction in the vegetative biomass of elite wheat plants likely contributes to the increasing seriousness of black-grass infestations in winter wheat crops. The reduction in root system size in wheat also makes elite wheat more prone to drought, because the plants are less able to access water stored at depth in the soil (Chapagain et al., 2014). Similarly the reduction in canopy size exaggerates the effect of fungal leaf infections in modern elite wheat, because there is so little ‘spare’ green leaf area if tissue is lost to infections. In essence, modern elite wheat is remarkably high-yielding under perfect conditions, but much more prone to losses in the face of biotic and abiotic stress, because its vegetative tissue has been pared back to the minimum.

Our results suggest that one way to breed winter wheat for increased competitiveness against black-grass (and other weeds) would be to focus on restoring root system size. Root traits have been shown to increase competitiveness against weeds in other crop species such as sugar beet (Stevanato et al., 2011) with root traits linked with increased nitrogen uptake being important in increasing weed suppression (Casper and Jackson 1997; Tilman and Wedin, 1991). While increasing root system size would require reversing the general direction of wheat breeding from recent years, there are a range of other current motives and opportunities that would support such a strategy. A move towards lower-input agriculture requires larger, more dense root systems better able to utilise nutrients, both the residue present in the soil, and those such as may still be applied as fertiliser. Increasingly erratic rainfall, and the increase in both drought and water-logging, require larger, deeper root systems to mitigate against these stresses. And the desire to increase storage of carbon within the soil also supports the breeding of crops with larger root systems (D)gnac et al., 2017). Increasing wheat root system size may also contribute to improving soil health (Chaparro et al., 2012). Thus, breeding for increased wheat root system size would create a more resilient, durable and sustainable crop, and would contribute to combatting black-grass infestations. Our results suggest that there is sufficient variation within current elite wheats to allow breeding for increased root system size (Figure 4), although introgressing variation from outside elite wheat may also be necessary to do this.

### Other routes to improving wheat competitiveness against black-grass

In the shorter-term, we have shown that there is some variation in competitive ability amongst elite wheat lines – irrespective of the root cause – which provides the possibility of providing farmers with a “recommended list” of varieties of both winter wheat and barley to grow in fields known to be infested with black-grass. These data could be integrated into the standard UK AHDB Recommended Lists or be provided as a supporting document for varietal selection. In line with the results of our landrace varieties that showed increased competitive ability compared to most elite varieties, older wheat varieties including the 1963 variety Park, were found to be more yield-stable under differing weed pressures than modern semi-dwarf cultivars (Mason, Goonewardene and Spaner, 2008). This highlights the presence of a treasure trove of competitive traits in older wheat varieties that may have been lost, but could be restored to elite wheat germplasm. The development of our relatively quick and easy screening method for crop - black-grass competition will hopefully also make it easier for future screening for competitive cultivars. Carrying out further field-scale trials with more crop lines will be vital to ensure the results of our competition screen are scalable to the field.

A non-mutually exclusive possibility to breeding for increased root system size may be to instead breed winter wheat for increased growth rate under winter conditions. Our results show that wheat grows slowly in winter conditions, but given the faster growth of barley and black-grass, this does not seem to be due to any environmental limitation. Thus, it seems very likely that increasing wheat is physiologically possible. Increasing winter growth rate in wheat would prevent black-grass gaining such a large competitive advantage, and would increase root system growth – with all the advantages outlined above – without necessarily changing the shoot:root ratio of the plant, and without having to specifically breed for increased root growth. Increasingly warmer winters may de-risk the strategy of increased winter growth rate, relative to colder winters of the past. It would also be interesting to know how variation in the early vigour of wheat lines affects their long-term ability to compete with black-grass. Bastiaans, Paolini and Baumann (2008) highlighted that early relative emergence improves access of the crop to resources, so would producing a larger shoot/root system prior to winter impact the crop’s ability to compete with black-grass and possibly negate the advantages accrued by black-grass over winter? Further investigation is also required into chemical exudate production by wheat and black-grass root systems, and whether this has any impact on the outcome of wheat – black-grass competition. It is certainly the case for instance that production of the allelochemical momilactone B influences the outcome of competition between rice and barnyard-grass (Minh and Xuan., 2024). It may be that black-grass inhibits wheat growth via allelochemicals, or that improving allelochemical production in wheat might help combat black-grass, but more research is needed to establish whether there is any chemical dimension in this particular crop-weed system. Overall, there appear to be range of options available to reduce the seriousness of black-grass infestations, while also helping to move towards lower-input agricultural systems with reduced dependence on fertilizer and herbicide applications.

## SUPPLEMENTARY DATA

**Supplemental Table 1: Experimental set-up for container trials**

## ACKNOWLEDGEMENTS

The authors acknowledge the support of the ADAS Boxworth field team in the set-up, running and harvesting of the container and field trials.

## AUTHOR CONTRIBUTIONS

J.C. performed experiments and analyzed the data. J.C, L.T and T.B. designed the study. All authors contributed to writing the manuscript.

## DATA AVAILABILITY

All figures in this manuscript are associated with raw data. All raw data will be made available upon request to the corresponding author.

